# Gradual Cerebral Hypoperfusion Impair Fear Conditioning and Object Recognition Learning and Memory in Mice: Potential Roles of Neurodegeneration and Cholinergic Dysfunction

**DOI:** 10.1101/177121

**Authors:** Jogender Mehla, Sean Lacoursiere, Emily Stuart, Robert J. McDonald, Majid H. Mohajerani

**Author notes:** Robert J McDonald and Majid H. Mohajerani are corresponding authors. Correspondence to or. **Correspondence: Dr. MajidH Mohajerani** Assistant professor, Department of Neuroscience, Canadian Centre for Behavioural Neuroscience, University of Lethbridge, 4401 University Dr West, Lethbridge-T1K 3M4, Alberta, Canada, **Dr. Robert J McDonald**, Professor, Department of Neuroscience, Canadian Centre for Behavioural Neuroscience, University of Lethbridge, 4401 University Dr West, Lethbridge-T1K 3M4, Alberta, Canada.

## Abstract

In the present study, male C57BL/6J mice were subjected to gradual cerebral hypoperfusion by implanting an ameroid constrictor (AC) on the left common carotid artery (CCA) and a stenosis on the right CCA. In the sham group, all surgical procedures were kept the same except no AC was implanted and stenosis was not performed. One month following the surgical procedures, fear conditioning and object recognition tests were conducted to evaluate learning and memory functions and motor functions were assessed using a balance beam test. At the experimental endpoint, mice were perfused and brains were collected for immunostaining and histology. Learning and memory as well as motor functions were significantly impaired in the hypoperfusion group. The immunoreactivity to choline acetyltransferase was decreased in dorsal striatum and basal forebrain of the hypoperfusion group indicating that cholinergic tone in these brain regions was compromised. In addition, an increased number of Fluoro-Jade positive neurons was also found in cerebral cortex, dorsal striatum and hippocampus indicating neurodegeneration in these brain regions. Based on this pattern of data, we argued that this mouse model would be a useful tool to investigate the therapeutic interventions for the treatment of vascular dementia. Additionally, this model could be employed to exploit the effect of microvascular occlusions on cognitive impairment in the absence and presence of Alzheimer pathology.

## INTRODUCTION

Vascular dementia (VaD) is the commonest form of dementia after Alzheimer’s disease which accounts for at least 20% of all causes of dementia [1]. Prevalence for VaD from developing countries has been reported in between 0.6 to 2.1% in those 65 years and older [2] with men more prone to VaD than women [4]. Interestingly, the incidence of VaD in the USA is around 3.8 per 1000 per annum [3]. VaD is caused by a reduction in cerebral blood flow by blockage of cerebral blood vessels which leads to impairments in cognitive function and is often associated with cardiovascular disease [5]. Continuous reduction in regional cerebral blood flow (CBF) exacerbates memory dysfunction and leads to the development and progression of dementia [7]. Gradual reduction in CBF due to arteriovenous malformations, carotid stenosis/occlusion, and cerebral small vessels disease are the leading causes of VaD [8-10]. Various diseases such as hypertension, diabetes and atherosclerosis are major risk factors for VaD [11]. Chronic reduction in CBF may lead to neurodegeneration via oxidative stress, neuroinflammation and impairment of neuronal energy system which may then cause cognitive impairments [7, 12]. Out of several forms of VaD, subcortical associated dementia is an area of research interest among neuroscientists because its high incidence and prevalence rate which affects daily living activities in individuals suffering from this disorder [6].

The chronic diffuse cerebral ischemia models are considered important for studying the correlation between chronic cerebral hypoperfusion and cognitive functions. Recently, Hattori et al. developed a potentially relevant mouse model of subcortical infarct lesions with cognitive impairment which may be used for development of novel therapeutic interventions for cerebral stroke and VaD [13]. However, we noted several limitations of the report using this model of VaD. First, only spatial learning and memory function was assessed. Since spatial learning tasks have a navigational component to them, a general motor impairment caused by the stroke could account for the learning and memory impairment. Accordingly, we selected tasks that had minimal motor requirements to provide stronger evidence for a clear learning and memory impairment in this model of VaD. Second, the Hattori et al did not evaluate the effects of VaD on cholinergic function [13]. This was deemed relevant because reduced cholinergic function is found in the aged and those that suffer from various forms of dementia [14]. Finally, we assessed the role of neurodegeneration. Therefore, to extend the research conducted by Hattori et al and to validate this as a mouse model of VaD, the present study was designed with assessment of non-spatial learning and function, cholinergic function and a measure of neurodegeneration.

## MATERIALS AND METHODS

### Animals

Male C57BL/6J mice (4-5 months old), weighing 25-30 g, were used in the present study. They were group housed 4 per cage and had food and water ad libitum. They were maintained on a 12:12 light/dark cycle. All experimental procedures were approved by the University of Lethbridge animal welfare committee and performed in accordance with the standards set out by the Canadian Council for Animal Care.

### Experimental groups

The mice were divided into two groups. Group 1 (n= 5) was designated the sham group where surgery was performed without implanting the ameroid constrictor (AC, Research Instruments NW, 30094 Ingram Rd, Lebanon, OR 97355, USA) on left common carotid artery (CCA) and without ligation of right CCA for stenosis. The inner diameter was 0.5 mm, the outer diameter 3.25 mm, and the length 1.28 mm. Group 2 (n= 8) was the gradual hypoperfusion group in which the AC was implanted on the left CCA as well as ligation of the right CCA for stenosis [13, 15]. The behavioral assessments were performed one month after surgery by an experienced researcher blinded to the experimental groups. After completion of the behavioral testing, the mice were perfused and brains were collected for histological and immunostaining assessments.

### Surgical procedure

For surgery, the mice were anesthetized with 1.5% isoflurane. Rectal temperature was maintained between 36.5 and 37.5°C. A midline cervical incision was given to expose both CCAs and they were separated from their sheaths. Two 6-0 silk sutures were placed around the distal and proximal segments of the left CCA. The artery was gently lifted by the sutures and placed in internal lumen of AC just below the carotid bifurcation. To ligate the right CCA, a 33G needle (external diameter = 0.20 mm) was positioned beside the CCA and tied together with a 6-0 silk suture; the needle was then quickly withdrawn. In the present study, we slightly changed the procedure for induction of sub-cortical infarcts as reported in previous study [13]. Here, we used a 33G needle instead of microcoil to make surgery simple. The midline incision was then sutured and the mice were transferred to a recovery room. Relative CBF was measured on presurgery day and on day 1, 3, 7, 14 & 28 postsurgery using laser speckle flowmetry, which has a linear relationship with absolute CBF values and obtains high spatial resolution 2D imaging as described in previous studies [16-18]. The recordings were performed through a glass cover slip cranial window under anesthesia with 1.0-1.2% isoflurane [19, 20]. The mean CBF was measured in identically sized regions of interest (located 2 mm lateral and 1 mm posterior from the bregma) using ImageJ as described by Mohajerani et al [16]. These reflectance optical signal reflect the CBF of the subsurface microvessels in the cortex [18]. CBF values were expressed as a percentage of the pre-surgery value. The subjects (n= 4) used for CBF were different from those used for behavioural assessment and histology in both sham and hypoperfused groups.

### Behavioral assessment

#### Object recognition test

The object recognition test was conducted as described by Vogel-Ciernia and Wood [21]. This test is based on the principle that mice made familiar with a specific object during training will spend more time exploring a novel object on the test day. For this task, mice are first trained to form object memories of two identical items [22]. A white plastic square box (52 x 51 x 30 cm) was used for object recognition training and testing. Two 100 ml beakers flipped upside down were used as familiar objects on training day and a small plastic box was used as the novel object.

Briefly, mice were brought from their home cage to the experimental room and placed in the testing box for 5 min daily for 6 days to habituate the subjects to the training environment and reduce the levels of fear/anxiety factors during the experiment. Then, 24 h after the habituation period, a training test was performed. The two familiar objects (beakers) were cleaned with 10% ethanol to mask any previous order cues and then allowed to dry completely. These objects were then placed in the testing box at equal distances from the walls in a central position. A video camera (30 frames/sec) was setup above the testing box to record the subjects’ behaviour for further analysis. The mice were introduced into the testing box for 10 min to explore both objects and video recordings were made for each mouse. The test session was performed 24 h after training in which one location contained the familiar object and the other position contained the novel object (plastic box). For the test, mice were individually placed in the testing box for 5 min to explore the objects and their behaviour was video recorded. After each mouse completed the test, feces were removed and bedding was stirred to equally distribute any odor cues. The objects were also wiped with 10% ethanol to mask the odor cues after each mouse. The data was analyzed by measuring the exploration time for familiar and novel objects during training and testing days. Then, an investigation ratio was calculated for each group.

#### Fear conditioning test

A fear conditioning test was performed as described previously with slight modifications [23]. This paradigm clear assessment of learning and memory functions supported by aversive associative conditioning processes. This test is used to assess the amygdala and hippocampus dependent memory in rodents. Fear conditioning was performed in acrylic square box (33 x 33 x 25 cm) located in a well-lit room. The front and rear walls consist of black acrylic material whereas white acrylic material was used for side walls. A video camera (30 frames/sec) was fixed to record the mice’s behavior for further analysis. The floor of chamber consisted of 64 stainless steel rods (2 mm diameter) spaced 5 mm apart. The rods were connected to a shock generator for the delivery of foot-shock. A speaker was used for delivery of the tone stimulus. Prior to conditioning, the chamber was cleaned with a 10% ethanol solution to mask any previous odor. The tone test was conducted in a chamber that was different from the conditioning chamber. This chamber was triangular and the walls consisted of white acrylic materials (33 x 33 x 29 cm). Prior to the tone test, the chamber was cleaned with a 10% ethanol solution.

On the conditioning day, mice were brought from their home cage into a testing room and allowed to sit undisturbed in their cages for 10 min. Mice were then placed in the square conditioning box and allowed to explore for 2 min before the onset of the tone (20 sec). In the delay conditioning procedure, a shock (2 sec, 0.5 mA) was given in the last 2 sec of tone presentations. The mice received five delayed conditioning trials, each separated by a 120-sec intertrial interval (ITI). The mice were taken out from the conditioning chambers 1 min after the last shock and returned to their home cages. 24 h later, the mice were placed in a different box than the conditioning box to assess conditioning to the tone in the absence of the training context. For the tone test, three 20-sec tones were presented after a 2-min baseline period. Each tone presentation was separated by a 120-sec ITI. The freezing response was measured using a time sampling procedure in which an observer scored the absence or presence of the freezing response for each animal at every 2 sec. Then 24 h after the tone test, the mice were placed back in the original conditioning box for 5-min context test. During this test, freezing of the animal was scored at every 5 sec. Data was transformed into a percent freezing score by dividing the number of freezing observations by the total number of observations and multiplying by 100. During the testing sessions, the boxes were cleaned with 10% ethanol after each mouse trial to mask any odor cues left by the previous subject.

#### Balance beam test

This task is used to assess sensorimotor function and balance in rodents. This test was performed as described previously with slight modifications [24]. Briefly, a 90 cm long metal beam with a round surface (diameter = 1 cm) was fixed 50 cm above a table top on two poles and a black escape box was placed at one end of the beam. The mice were placed at the other end of the beam and the time required to cross to the escape box at the other end of beam was recorded. The mice were given 3 training sessions to learn how to walk on the beam. After the training session, the mice were returned to their home cage. 24 hr after the training, testing was performed by giving more 3 trials to each mouse. Time to cross the beam and the number of paw slips from the beam were recorded. A paw slip was defined as a foot coming off the top of the beam. The data from 3 testing trials was averaged. The beam and escape box were cleaned with 10% ethanol after each trial.

### Fluoro-Jade staining

Fluoro-Jade staining was used as a marker of damaged neurons [25] and was performed as described previously with some modification [26]. We analyzed 3 brain sections for sensorimotor cortex (+0.26-0.50 mm from bregma; 0.6 mm^2^ each section located between 2-3 mm from midline ± 0.5 mm) from each mouse. The mice brains were sectioned (thickness-40 μm) using cryostat. Sections were then mounted on gelatin coated slides and dried at room temperature overnight. The next day, sections were rinsed in distilled water for 1 min. The sections were incubated in freshly prepared 0.06% potassium permanganate for 15 min on an orbital shaker and washed with distilled water for 2 min. The sections were then stained with 0.0004% Fluoro-Jade B in 0.1% acetic acid for 30 min. After 3 washes in distilled water, the sections were coverslipped. Lastly, stained sections were imaged under 10x & 20x magnification using laser scanning confocal microscope (Olympus FLUOVIEW FV1000, Olympus, JAPAN). The numbers of neuron stained positive for Fluoro-Jade were analyzed using Image-J software.

### Choline acetyltransferase (ChAT) immunostaining

The immunohistochemistry for ChAT was done as described previously with some modification [27]. We analyzed 3 sections from MSDB complex (+0.74-1.18 mm from bregma; 84x142 mm2 each section located between 0-0.6 mm from midline) for each mouse. Floating sections (40 μm thick) were washed in PBS for 10 min and blocked in a solution containing 5% normal goat serum and 0.1%TritonX-100 for 30 min. Sections were then incubated at 4°C with anti-ChAT primary antibody (monoclonal rabbit, Abcam, ab178850, 1:5000) overnight. Then three washes (10 min each) were given and sections were again incubated with Anti-rabbit-Alexa-594 secondary antibody (Anti-rabbit IgG (H+L) goat, invitrogen-Life Technology A11037, 1:500) for 4 hrs. The sections were mounted on microscope slides, counterstained with DAPI, dehydrated, and covered with coverslips. Finally, the sections were imaged under 40x using Nanzomer microscope (NanoZoomer 2.0-RS, HAMAMTSU, JAPAN). The numbers of neuron stained positive for ChAT were analyzed using Image-J software.

### Statistical analysis

The statistical analysis was performed using SPSS statistical software package version 22.0. Results were presented as mean ± SEM. The statistical significance difference between the groups was evaluated using Student’s t-test. CBF data was also analyzed using Student’s t-test. A p value < 0.05 was considered as statistically significant.

## RESULTS

In the present study, we did not observed any mortality neither in sham operated group nor in surgery operated group. A significant gradual decrease in CBF was found from day 1 to day 7 and maximum reduction CBF was observed on day 7. Later, between the period of the 14^th^ and 28^th^ day, an improvement of CBF was observed (Fig. 1A). No significant change in CBF was found in sham group as compared with presurgery day. Additionally, hypoperfused group showed the significant reduction in CBF when compared with sham group (Fig. 1A).

**Fig. 1.**
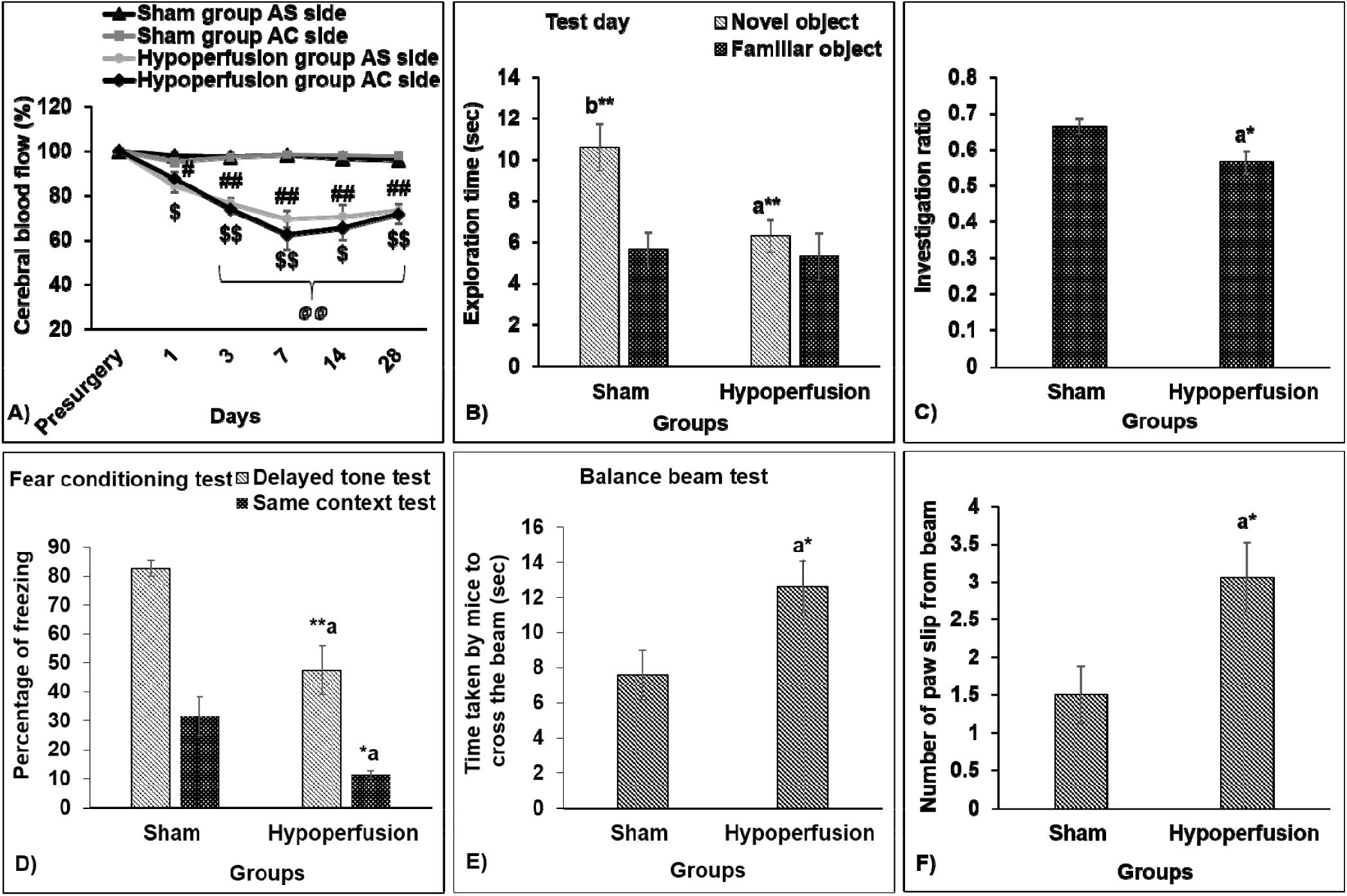
A) Measurement of cortical surface cerebral blood flow (CBF) using laser speckle flowmetry (N=4 for each group). Evaluation of learning, memory and motor performance of mice using an object recognition test, fear conditioning test and balance beam test, respectively. In the object recognition test, memory in mice was evaluated by measuring: B) exploration times during test day; C) investigation ratio. In the fear conditioning test, memory in mice was evaluated by measuring: D) percentage of freezing. In the balance beam test, motor performance was evaluated by measuring: E) time taken by mice to cross the beam; F) number of paw slips from beam. All values are presented as mean ± SEM. *p<0.05, **p<0.01; a-as compared to compared to sham group; b-as compared to familiar object exploration time of sham group. ^#^p<0.05, ^##^p<0.01 as compared to pre-surgery CBF for artery stenosis (AS) side; ^$^p<0.05, ^$$^p<0.01 as compared to pre-surgery CBF for ameroid constrictor (AC) side; ^@@^p<0.01 as compared to sham group. N= 5 in sham group; N= 8 in hypoperfusion group. AS side stands for artery stenosis side; AC side stands for ameroid constrictor side

In the novel object recognition test, 24 hr after the acquisition training, when mice were placed back in the testing box with one similar object replaced with a novel object, the hypoperfused mice spent significantly (t(11)= 3.181, p= 0.009) less time exploring the novel object as compared to sham mice (Fig. 1B). On test day, sham mice showed significantly (t(4)= 7.003, p= 0.002) longer exploration time for the novel object in comparison to the familiar object (Fig. 1B) whereas no significant (t(7)= 1.086, p= 0.313) difference was found between exploration time of novel and familiar objects in the hypoperfused group (Fig. 1B). Moreover, the investigation ratio for sham mice was also significantly (t(11)= 2.262, p= 0.045) higher than the hypoperfused mice (Fig. 1C). The results of this experiment suggest that chronic hypoperfusion reduces the mice ability to distinguish a new object from one that has been encountered previously.

In the delayed tone fear conditioning test, the hypoperfused mice group showed significantly (t_(11)_= 3.959, p=0.002) less percentage of freezing as compared to sham mice group (Fig. 1D). Moreover, when hypoperfused mice were again placed in the original square training box for 5 min 24 hr after the delayed tone test, these mice also showed significantly (t_(11)_= 3.011, p= 0.012) lower contextual freezing levels in comparison to sham mice (Fig. 1(D)). These results suggest that learning and memory functions associated with aversive classical conditioning to cues and contexts was impaired in the hypoperfused mice.

The results from the balance beam test indicated an impairment of motor function in the hypoperfused mice as well. The hypoperfused animals took significantly (t(11)= -2.296, p= 0.0.042) more time to cross the beam as compared to sham mice (Fig. 1E). Moreover, the number of paw slips from the beam were also significantly (t(7)= -2.365, p= 0.037) higher in the hypoperfused mice in comparison to sham mice (Fig. 1F).

For histology/immunohistochemistry, we analyzed 3 brain sections from each mouse and analysis was conducted by researcher blinded to experimental groups. We found an increase in the number of Fluoro-Jade positive neurons in the constricted side of the cerebral cortex (t_(4)_= - 9.710, p= 0.001), dorsal striatum (t_(11)_= -14.038, p= 0.000) and CA1 of hippocampus (t_(11)_= - 13.742, p= 0.000) of the hypoperfused group in comparison to the sham group (Fig. 2B, D; 2F, H; 2J, L; see panel M-O for the summary results; n = 5 mice per group). Moreover, the hypoperfused group also showed a significant increase in the number of Fluoro-Jade positive neurons in the constricted side of the cerebral cortex (t_(2)_= -9.693, p= 0.01), dorsal striatum (t_(2)_= -15.457, p= 0.004) and CA1 of hippocampus (t_(11)_= -13.578, p= 0.005) in comparison to the ligated side (Fig. 2C, D; 2G, H; 2K, L; see panel M-O for the summary results; n = 5 mice per group). We also observed a significantly increased number of Fluoro-Jade positive neurons in dorsal striatum (t_(4)_= -5.205, p= 0.006) and CA1 of hippocampus (t_(11)_= -4.630, p= 0.01) in the ligated side of the hypoperfusion group in comparison to sham group (Fig. 2E, G; 2I, K; see M-O for summary results; n = 5 mice per group).

**Fig. 2.**
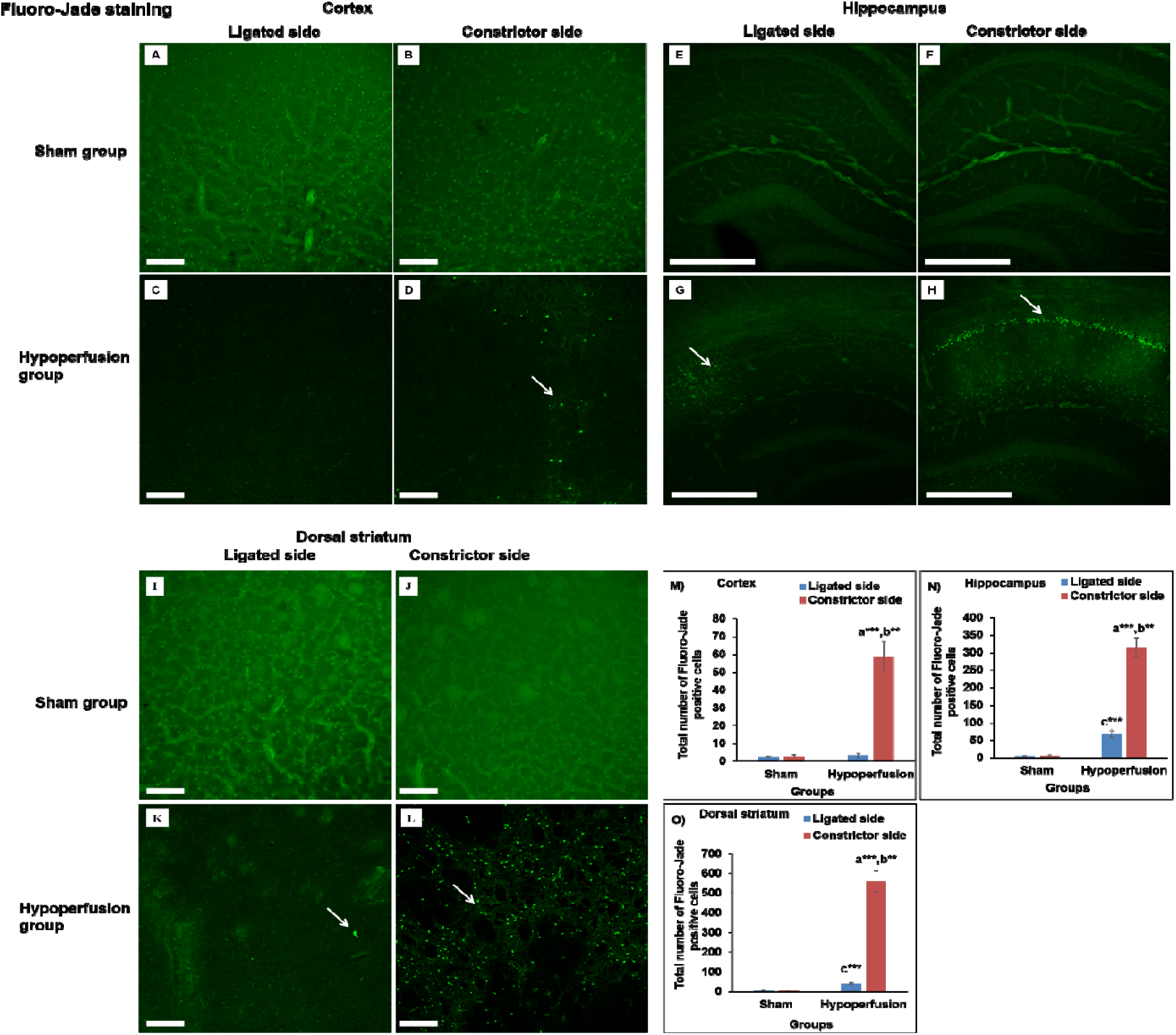
Photomicrographs of Fluoro-Jade staining [cerebral cortex (A,B,C,D; magnification- 20x), hippocampus (E,F,G,H magnification-10x)] and dorsal striatum (I,J,K,L; magnification- 20x) of sham and hypoperfusion mice. The hypoperfusion mice showed an increased number of Fluoro-Jade positive cells in the cerebral cortex, dorsal striatum and hippocampus on the constricted side and some in dorsal striatum and hippocampus of ligated side. Scale bar-100 μm for cerebral cortex; Scale bar-250 μm for hippocampus. Data are presented as mean ± SEM. **p<0.01, ***p<0.001; a-as compared to compared to sham group; b-as compared to ligated side of hypoperfusion group; c-as compared to ligated side of sham group. N= 5 mice in each group.

The ChAT positive cells in the hypoperfused group (65.2 ± 6.85) were decreased significantly (t_(4)_= 6.966, p= 0.002) in medial septal-diagonal band (MSDB) areas as compared to sham group (148.8 ± 7.05; Fig. 3G). Moreover, a significant (t_(4)_= 3.894, p= 0.018) decrease in ChAT positive neurons was also observed in the constricted side (47.0 ± 4.47) of dorsal striatum of the hypoperfused group as compared to the sham group (71.2 ± 6.15; Fig. 3H). Nonetheless, in comparison to ligated side (70.8 ± 6.79), the constricted side (47.0 ± 4.47) of dorsal striatum of the hypoperfusion mice showed significant (t_(2)_= 4.956, p= 0.038) less number of ChAT positive cells (Fig. 3 H).

**Fig. 3.**
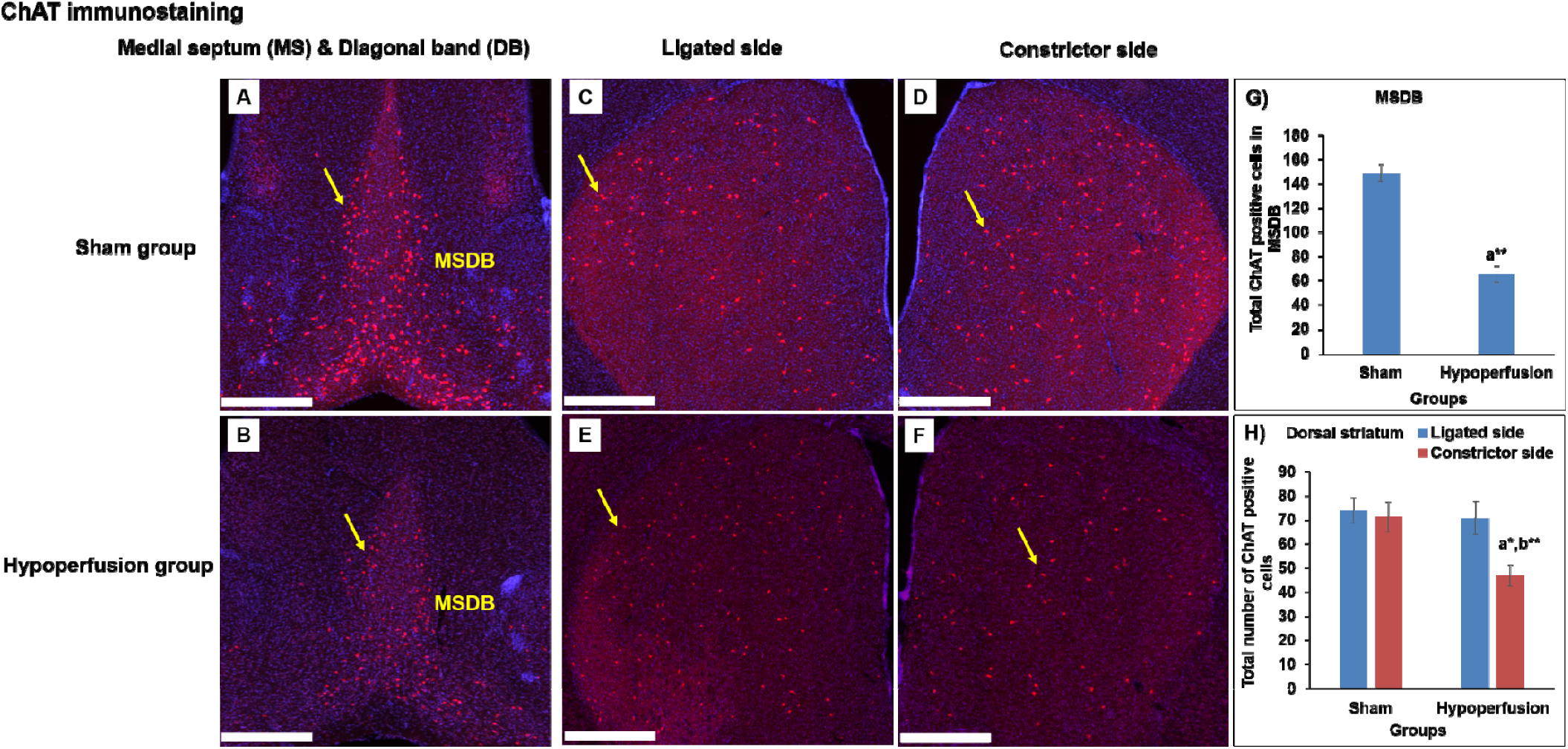
Photomicrographs of immunohistochemistry staining of choline acetyl transferase (ChAT) positive neurons in medial septum-diagonal band complex of the basal forebrain and dorsal striatum of sham and hypoperfusion mice. The hypoperfusion mice also showed decreased ChAT positive neurons in medial septum-diagonal band complex of the basal forebrain and dorsal striatum in comparison to sham group. Scale bar-500 μm, Magnification-40x. Data are presented as mean ± SEM. *p<0.05, **p<0.01; a-as compared to compared to sham group; b-as compared to ligated side of hypoperfusion group. N= 5 mice in each group

## DISCUSSION

Hattori et al reported a dementia-like condition in mice that experienced a specific gradual occlusion hypoperfusion which is more closely related to human vascular disease pathology than other experimental models of cerebral ischemia [28-30]. In that study, the investigators provided a good description of the damage induced by the gradual occlusion clearly indicating infarcts in subcortical areas including corpus callosum, internal capsule, hippocampus, fimbria, and caudoputamen [13].

However, we noted several limitations to that initial report. First, they only studied the functional impacts of the brain damage on a spatial version of the water task thought to be dependent on the hippocampus and related circuits. Therefore, to strengthen the validity and applicability of this novel model in vascular dementia related research, we proposed in the present study to evaluate non-spatial memory function using novel object recognition and fear conditioning tests. The former is sensitive to perirhinal cortex [22] and the latter is dependent on the amygdala [31] and hippocampal dysfunction [31].

The results showed that in cued, as well as context fear conditioning, the hypoperfused mice showed significantly less conditioned freezing behavior as compared to the sham mice, indicating learning and memory impairments previously shown to be dependent on neural circuits centered on the amygdala and/or hippocampus, respectively [31]. Additionally, the hypoperfused mice showed memory impairments on the novel object recognition test indicating dysfunction of cortical association circuits such as perirhinal and medial prefrontal cortices [22]. This pattern of impairments is in line with a previous study in which impairment of amygdala-associated memory was reported in a rat model of cerebral hypoperfusion [32] and a study by Yoshizaki and colleagues who reported memory impairments of mice in object recognition test in an experimental model of right unilateral CCA occlusion [30]. We also found an impairment of motor dysfunction on the balance beam task in the hypoperfused mice as compared to the sham group which is consistent with a previous studies in which impairments of motor function has also been reported in experimental model of cerebral hypoperfusion [13, 33, 34].

In addition to the functional effects of hypoperfusion described above, we found neurodegeneration in the cerebral cortex, dorsal striatum and hippocampus which may be responsible for the cognitive impairments we found in the hypoperfusion group. These neurodegeneration effects are in agreement with previous findings where neuronal loss in hippocampus has also been reported [13, 35-37].

The connection between cholinergic function and vascular dementia, learning and memory is well established in previous work [38, 39]. In the present study, we found decreased ChAT positive cholinergic neurons in the MSDB complex of basal forebrain and the dorsal striatum of the hypoperfused mice. These results are consistent with previous studies showing reduced levels of the ChAT using a bilateral common carotid artery occlusion model of cerebral hypoperfusion in rats [40-42]. These brain changes might contribute to the impairment in learning and memory function found in these rodent stroke models. Importantly, in the present study, the pattern of CBF reduction was similar to previous study [13].

In summary, the findings of the present study of the effects of gradual hypoperfusion induced subcortical infarcts showed clear impairments of learning and memory functions dependent on the amygdala, hippocampus and perirhinal cortex. These cognitive impairments were associated with neurodegeneration and cholinergic dysfunction. Since cerebral hypoperfusion, along with microvascular dysfunction and hypometabolism, in AD patients often occurs prior to cognitive dysfunction, it is possible that treatment of cerebrovascular changes and cerebral blood flow could be targeted to delay, or prevent, the progression of AD [43, 44]. Importantly, this experimental model could be used to understand the mechanisms of brain dysfunction and cognitive impairments found in subcortical induced vascular dementia and to assess the efficacy of novel drugs to prevent or treat this neurological disorder.

## ACKNOWLEDEMENTS

This work was supported by Natural Sciences and Engineering Research Council of Canada (NSERC) Discovery Grant #40352 and #06347 to MHM and RJM respectively, Campus Alberta for Innovation Program Chair (MHM), Alberta Alzheimer Research Program (MHM & RJM), Alzheimer Society of Canada (MHM & RJM), NSERC CREATE graduate scholarship (SL) and NSERC undergraduate student research award (ES). We thank Di Shao and Behroo Mirza Agha for animal breeding.

## AUTHOR CONTRIBUTIONS

JM performed the experiments and analyzed the results, SL helped with the experiments and analyzing the results, ES assisted in interpreting the results, MHM & RJM designed and supervised the experiments. JM, RJM & MHM contributed to the writing of the manuscript.

